# Systemic inflammation and risk of age-associated diseases in people living with HIV on long term suppressive antiretroviral therapy

**DOI:** 10.1101/418012

**Authors:** Hemalatha Babu, Anoop T Ambikan, Erin E Gabriel, Sara Svensson Akusjärvi, Alangudi Natarajan Palaniapan, Vijila Sundaraj, Naveen Reddy Mupanni, Maike Sperk, Narayanaiah Cheedarla, Rathinam Sridhar, Srikanth P Tripathy, Piotr Nowak, Luke Elizabeth Hanna, Ujjwal Neogi

**Affiliations:** Department of HIV/AIDS, National Institute for Research in Tuberculosis, ICMR, Chennai, India; Division of Clinical Microbiology, Department of Laboratory Medicine, Karolinska Institutet, Stockholm, Sweden; Department of Medical Epidemiology and Biostatistics and Institute of Environmental Medicine both of Karolinska Institutet Stockholm, Sweden; Government Hospital of Thoracic Medicine, Tambaram Sanatorium, Chennai, India; Department of Medicine Huddinge, Unit of Infectious Diseases, Karolinska Institutet, Karolinska University Hospital, Stockholm, Sweden; Department of Clinic, National Institute for Research in Tuberculosis, ICMR, Chennai, India

## Abstract

The ART program in low- and middle-income countries (LMIC) like India, has a public health approach with the standardized regimen for all people living with HIV (PLHIV). Based on the evidence from high-income countries (HIC), the successful implication and scale-up of ART in India, the risk of an enhanced and accentuated onset of premature-aging or age-related diseases could be observed in PLHIV. However, very limited data is available on residual inflammation and immune activation in the populations who are on first-generation anti-HIV drugs like zidovudine and lamivudine that had more toxic side effects. Therefore, the aim of the present study was to evaluate the levels of systemic inflammation and understand the risk of age-associated diseases in PLHIV on long-term suppressive ART using a large number of biomarkers of the inflammation and immune activation. Blood samples were obtained from therapy naïve PLHIV (Pre-ART, n=43), PLHIV on ART for >5 years (ART, n=53), and HIV-negative healthy controls (HIVNC, n=41). Samples were analyzed for 92 markers of inflammation, sCD14, sCD163, and telomere length. Several statistical tests were performed to compare the groups under study. Multivariate linear regression was used to investigate the associations. Despite a median duration of eight years of successful ART, sCD14 (p<0.001) and sCD163 (p=0.04) levels continued to be significantly elevated in ART group as compared to HIVNC. Eleven inflammatory markers, including 4E-BP1, ADA, CCL23, CD5, CD8A, CST5, MMP1, NT3, SLAMF1, TRAIL and TRANCE, were found to be significantly different (p<0.05) between the groups. Many of these markers are associated with age-related co-morbidities including cardiovascular disease, neurocognitive decline and some of these markers are being reported for the first time in the context of HIV-induced inflammation. Linear regression analysis showed a significant negative association between HIV-1-positivity and telomere length (p<0.0001). In ART-group CXCL1 (p=0.048) and TGF-α (p=0.026) have a significant association with increased telomere length and IL-10RA was significantly associated with decreased telomere length (p=0.042). This observation warrants further mechanistic studies to generate evidence to highlight the need for enhanced treatment monitoring and special interventions in HIV-infected individuals.

## Introduction

The most remarkable achievement in the battle against the Human Immunodeficiency Virus (HIV) is the discovery of very efficient, well-tolerated combinational antiretroviral therapy (ART), that has transformed the deadly viral infection into a chronic, manageable disease. In the absence of a cure, HIV-infection requires lifelong treatment. Though treatment successfully controls HIV-replication and prevents opportunistic infections, HIV-infected persons on long term ART not only succumb to death at an earlier age than the HIV-uninfected counterparts but also suffer from some maladies that are typically associated with human aging (Deeks, 2011). The effect of HIV-associated inflammation and immune activation on premature onset of immunosenescence despite effective viral suppression are thought to be the primary reasons for early aging in people living with HIV (PLHIV) (Sokoya et al., 2017).

Chronic, low-grade systemic inflammation resulting from an increased pro-inflammatory state contributes to the progressive pathophysiological changes associated with aging. This process of increase in pro-inflammation followed by a chronic inflammatory state is termed as “inflamm-aging” and is a significant risk factor for morbidity and mortality in the elderly people (Franceschi et al., 2000). The inflammatory environment triggers the development of several age-related noninfectious comorbidities (NICMs) (Deeks, 2011). In HIV infection, it is believed that viral persistence in a rare population of long-lived, latently infected cells despite successful ART contributes to the chronic inflammatory state (Deeks, 2011).

Unlike in high-income countries (HIC), the ART program in low-and middle-income countries (LMIC) like India, follows a public health approach with a standardized regimen for all PLHIV. As of December 2016, nearly one million PLHIV were receiving free ART through the government program in the country (UNAIDS, 2016). The recommended first-line ART regimen includes one non-nucleoside reverse transcriptase inhibitor (NNRTI), either nevirapine (NVP) or efavirenz (EFV), in the backbone of two nucleoside reverse transcriptase inhibitors (NRTI), either zidovudine (AZT) or tenofovir (TDF), and lamivudine (3TC). Although perfect adherence to treatment remains a challenge, the reasonably good response to first-line therapy indicates the overall success of the ART program in the country (Neogi et al., 2013). Thus, with the expansion of the ART program and its consequences in PLHIV, the burden of age-related non-AIDS diseases are likely to increase. Since the environment has an enormous impact on age and age-related diseases, and the genetic determinants of aging may vary across populations, studies conducted in HICs might not apply to the LMICs (Hoffman et al., 2017).

The present study attempted to evaluate HIV-associated inflammation and immune activation and the contribution to inflamm-aging in PLHIV on long-term combination ART (cART) by assessing markers of inflammation as well as aging, including a panel of 92 inflammatory markers, two well-characterized immune activation markers (sCD14 and sCD163), and telomere length. The study provides useful insights into the role of inflammation and aging in HIV-1 infected individuals despite successful ART.

## Materials and Methods

### Study design and participants

The study included three groups of individuals: i) treatment-naïve PLHIV with viremia and moderate CD4 count (Pre-ART herein) ii) PLHIV of aged between 35-60 years on ART for more than five years with suppressed viremia on national first-line ART and >90% adherence to treatment (ART herein), and iii) life-style, age and gender-matched (with ART group) HIV-1 negative healthy individuals free of any kind of chronic illness (HIVNC herein). The HIV-1 positive cohort was recruited from a tertiary care ART Centre at the Government Hospital for Thoracic Medicine (GHTM), Chennai, India, at the time of routine standard-of-care hospital visit. Exclusion criteria were pregnancy in women, Immune Reconstitution Inflammatory Syndrome (IRIS), presence of co-infections like active tuberculosis or hepatitis virus infection, history of co-morbidities like diabetes mellitus, obesity, evidence of cardiovascular disease or any chronic illness, illicit drug usage, alcohol consumption, and intake of anti-inflammatory drugs in the past one month. For the ART group, we screened 258 individuals, and recruited 55 persons who matched the above criteria for the study. After screening 166 individuals for the pre-ART group, 41 who met the eligibility criteria were recruited for the study. Plasma viral load was measured using the *Abbott* RealTime HIV-1 viral load assay (Abbott, US). Two individuals in the ART group had viral load >150 copies/mL (4000 and 1800 copies/mL, respectively) and had to be excluded from the study. We screened 295 healthy individuals in and around Chennai, India, and identified 43 life-style, age and gender-matched individuals for the HIVNC group.

The overall study design is presented in Figure 1. After first-time counseling and obtaining informed consent to participate in the study, 15 mL of venous blood was collected from each study participant.

**Figure 1.**
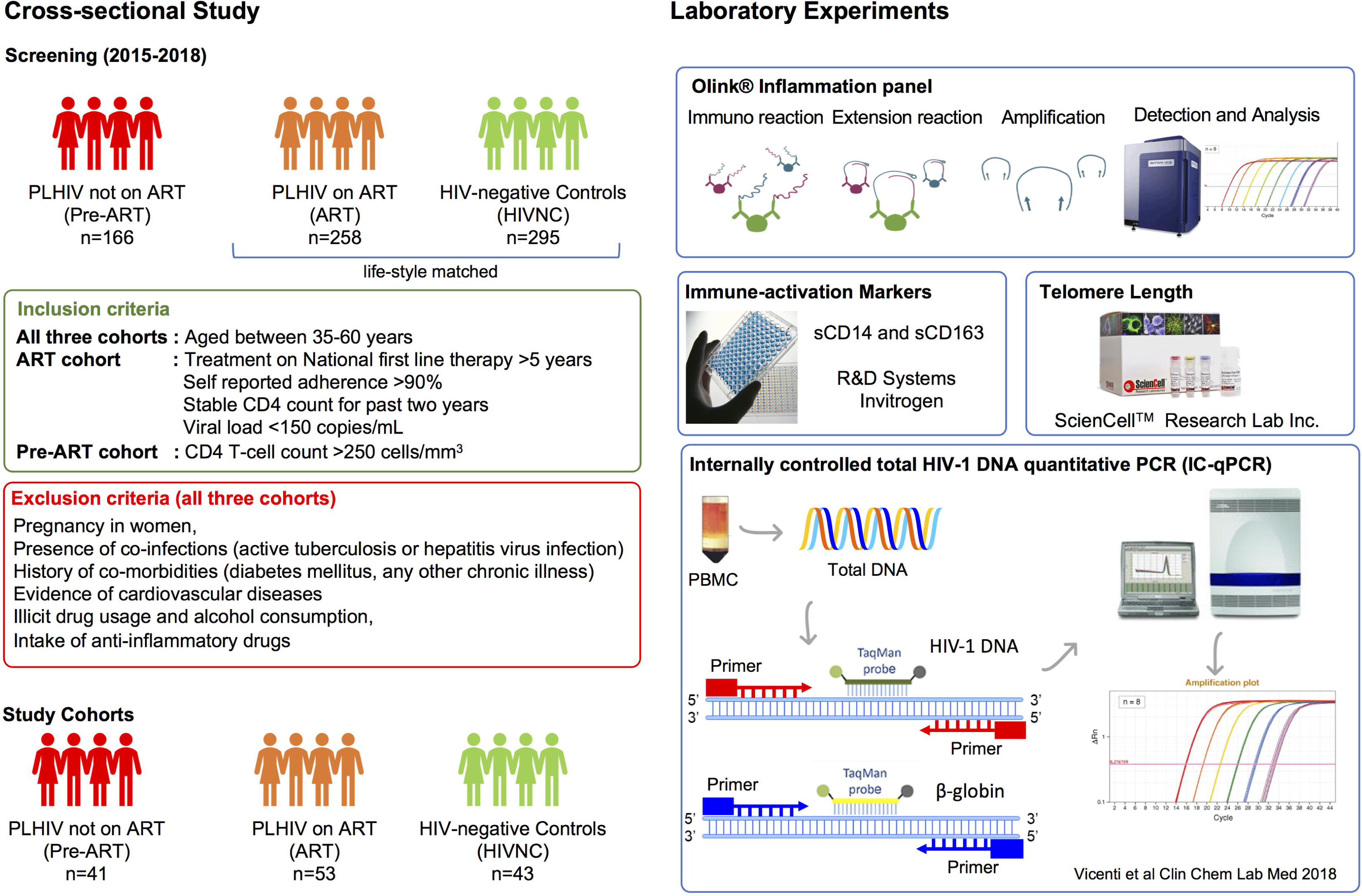
Flow diagram of study design and experimental plan: 424 HIV-1 positive individuals and 295 HIV-1 negative healthy controls were screened. Following defined inclusion and exclusion criteria, 43 healthy controls, and 53 HIV positive ART-experienced subjects and 41 ART-naïve HIV-1 positive subject were recruited for the study. The methodology used for the study was also presented.

### Proteomic profiling of the soluble factors in plasma

Plasma samples from the Pre-ART, ART and HIVNC groups were subjected to soluble proteome analysis using the Olink^®^ Inflammation Panel that includes 92 inflammation-related soluble factors (Olink Bioscience AB, Uppsala, Sweden) (Assarsson et al., 2014). We measured 92 markers of inflammation in the plasma samples of the three study groups. Of the 92 proteins, only 75 were detectable in >50% of the samples, and therefore our analysis was restricted to these proteins alone. Two Pre-ART samples and one HIVNC sample did not pass the quality control and had to be excluded from our analysis. We also measured two widely reported biomarkers of immune activation, sCD14 (Human CD14 Quantikine ELISA Kit R&D Systems, UK) and sCD163 (Thermo Scientific^™^ Pierce^™^ Human CD163 Kit, Thermo Scientific, USA) in plasma.

### Telomere length in peripheral blood mononuclear cells (PBMC)

As a molecular biomarker of “biological aging”, the leucocyte telomere length was measured in PLHIV on ART and HIV-negative controls. Genomic DNA was extracted from PBMC using the QIAamp *DNA Mini Kit (Qiagen, Germany).* and telomere length was measured using the Absolute Human Telomere Length Quantification qPCRAssay Kit (AHTLQ; ScienCell Research Laboratories, US) following manufacturer’s instructions. All samples were tested in triplicate.

### Total HIV-1 DNA quantification using IC-qPCR marker for HIV-1 reservoir level

To quantify total HIV-1 DNA from PBMC, internally controlled qPCR (IC-qPCR) was performed as described previously (Vicenti et al., 2018). IC-qPCR was performed in duplicate using 500ng template DNA and Takara Premix Ex Taq™ (Probe qPCR) (Takara, Japan). Primers targeting HIV-1 LTR and Beta-globulin were used. Total HIV-1 DNA copy numbers was calculated based on the linear equation of the 10-fold Beta-globulin standard curve derived from Jurkat cells and the 10-fold pNL4-3 plasmid standard curve, diluted in 50 ng/μL of Jurkat DNA to mimic clinical samples, and normalized to obtain HIV-1 DNA copies per million PBMC.

### Statistical analysis and data visualization

Mann Whitney U test, Chi-square test, and one-way analysis of variance (ANOVA) were performed to investigate difference in mean levels of protein expression (NPx) in the different groups under study. Multivariate linear regression for the outcome of telomere length was used to investigate the association between HIV status and HIV treatment duration adjusting for chronological age as well as other disease and patient characteristics. These same models were used to investigate evidence of mediation of the HIV-telomere relationship by soluble biomarkers. A heatmap was generated to visualize the clustering of samples based on protein expression using *gplots* v3.0.1 in R. Similarity in protein expression within the groups was visualized using a multi-dimensional scaling plot (MDS) in R package, edge R. All other analysis was performed using base R. P-values are not corrected for multiple comparisons, although false discovery rate was used in some of the biomarker discovery analyses.

### Ethical Clearances

The study was approved by the Institutional Ethics Committee of the National Institute for Research in Tuberculosis (NIRT IEC No: 2015023) and Institutional Review Board of the Government Hospital for Thoracic Medicine (GHTM-27102015) Chennai, India. All the study participants gave written informed consent. Patient identities were anonymized and delinked before analysis.

## Results

### Patient characteristics

Cohort characteristics are presented in Table 1. All three cohorts were gender matched. There were 51%, 43% and 51% of females in the Pre-ART, ART and HIVNC respectively. The median age of ART, Pre-ART and HIVNC groups were 45, 40 and 46 years respectively. In the ART group, the median (IQR) duration of treatment was 8 years (6-10 years). As per the inclusion criteria, all the patients had >90% of self-reported adherence which was further confirmed by viral load <150 copies/mL and very low-level HIV-1 reservoir with median (25^th^ −75^th^) 2.87 (2.62-3.18) log_10_ copies/10^6^ PBMC. Within the ART group, 57% (30/53) were on a zidovudine, lamivudine and nevirapine (ZDV/3TC/NVP) regimen and the remaining 43% (23/53) were on a tenofovir, lamivudine, and efavirenz (TDF/3TC/EFV) regimen. All individuals were initiated on antiretroviral treatment in the chronic phase of disease with a median (25^th^ −75^th^) CD4 count of 186 (100-280) cells/µL, as per the National guidelines for eligibility to ART existing at that point in time.

**Table 1.** Patients’ demographic and clinical parameter

| Parameter | Treatment naïve (Pre-ART) | Treatment Experience (ART) | HIV-negative Control (HIVNC) | P values |
| --- | --- | --- | --- | --- |
| N | 41 | 53 | 43 | ND |
| Gender, Female, N (%) | 21 (51) | 23 (43) | 22 (51) | 0.6734<sup>#</sup> |
| At sampling |  |  |  |  |
| Age in years, median (IQR) | 40 (37-43) | 45 (42-49) | 46 (40-54) | <0.0001* |
| CD4 count (cells/μL); median (IQR) | 367 (251-578) | 667 (476-797) | NA | <0.0001<sup>§</sup> |
| CD8 count (cells/μL); median (IQR) | 1138 (872-1625) | 772 (337-1092) | NA | <0.0001<sup>§</sup> |
| CD4:CD8 ratio, median (IQR) | 0.329 (0.1863-0.529) | 0.76 (0.575-1.013) | NA | <0.0001<sup>§</sup> |
| Viral Load, Log<sub>10</sub> copies/mL, mean (SD) | 4.4943 (0.9036) | 2.14 | NA | <0.0001<sup>§</sup> |
| Years on treatment, median (IQR) | NA | 8 (6-10) | NA | ND |
| Treatment Regimen, n (%)<br>ZDV+3TC+NVP<br>TDF+3TC+EFV | NA | 30 (57%)<br>23 (43%) | NA | ND |
| CD4 Count at treatment initiation (cells/μL), median (IQR) | NA | 186 (100-280) | NA | ND |
| HIV-1 Reservoir by total DNA count, log<sub>10</sub> copies/mL/10<sup>6</sup> PBMC, Median (IQR) | ND | 2.87 (2.62-3.18) | ND | ND |

NA: Not available, ND: Not Done, \*Kruskal-Wallis test, <sup>#</sup> $\chi^2$ test, <sup>§</sup>Mann-Whitney test
ZDV: zidovudine, 3TC: lamivudine, NVP: nevirapine, TDF: tenofovir and EFV: efavirenz, PBMC: Peripheral blood mononuclear cells.

### Soluble markers of immune activation in plasma

Soluble markers of immune activation, sCD14 and sCD163, were measured in plasma samples of the three study groups. PLHIV had significantly higher levels of sCD14 (**Figure 2a**) and sCD163 (**Figure 2b**) in plasma as compared to HIVNC (p<0.001, Mann Whitney U Test). Interestingly, despite prolonged suppressive ART (median duration of eight years of cART), there was no significant decline in the levels of sCD14 in PLHIV. Levels of sCD163 were significantly lower in the ART group as compared to the Pre-ART group (30014 pg/mL vs. 68192 pg/mL, p<0.001, Mann Whitney U Test), but were significantly higher as compared to the HIVNC group (p=0.04, Mann Whitney U Test). We did not find any significant correlation between duration of cART and sCD14 (Spearman r: 0.163; p=0.2432) or sCD163 (Spearman r: 0.154; p=0.2720) levels in the ART group.

**Figure 2.**
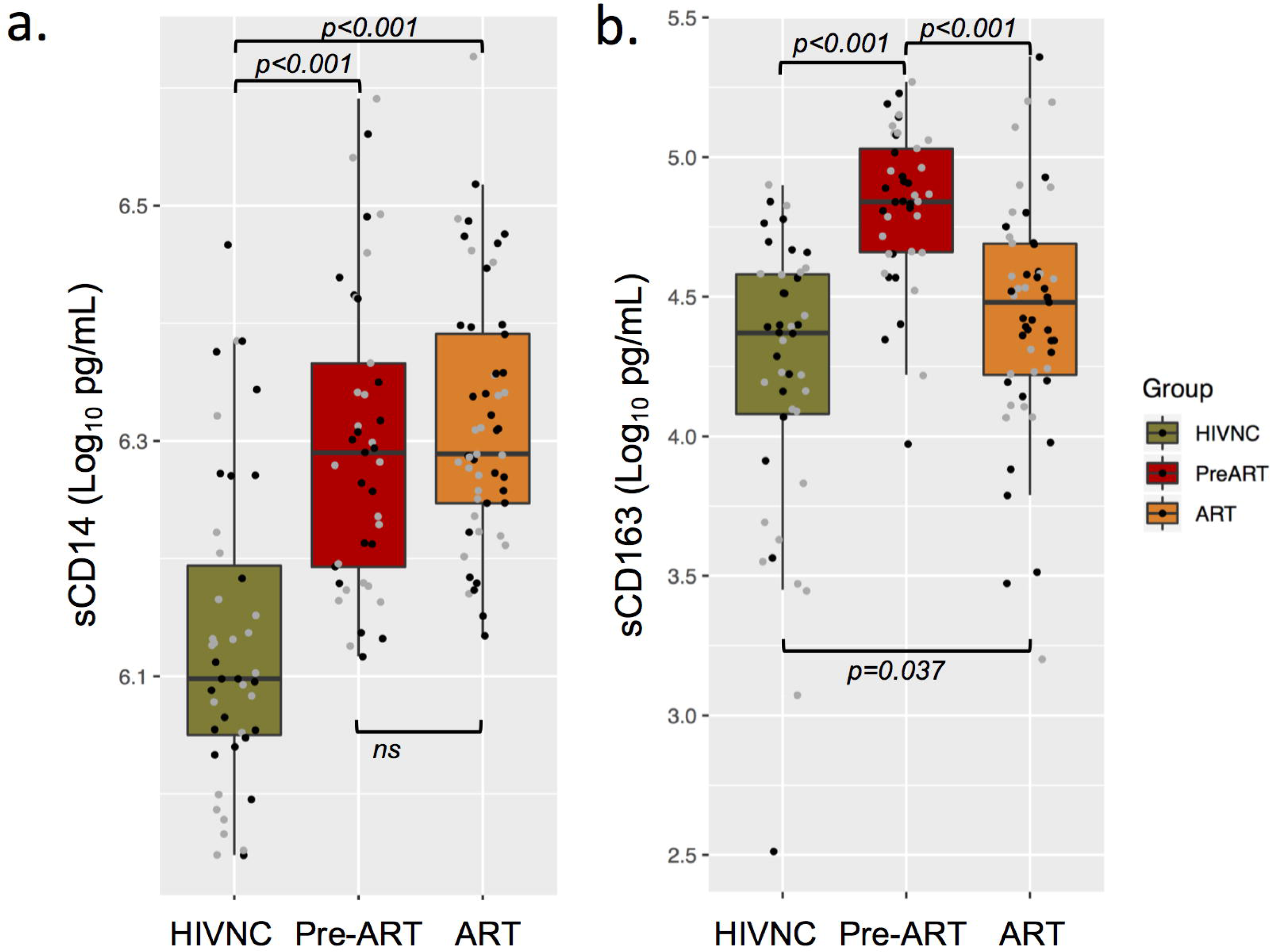
Plasma immune activation markers: Soluble CD14 (a) and CD163 (b) in plasma of the three groups of individuals were measured using ELISA.

### Soluble markers of inflammation in plasma

To identify candidate biomarkers for future study, we looked for biomarkers that differed significantly between the groups by Random Forest (RF) analysis (**Figure 3a**). The proteins with the most significant differences were TNFRSF9, sCD6, sCD5, TRANCE, and CXCL9. Hierarchical clustering analysis (HCA) with false discovery rate (FDR) <0.001, revealed clustering of 79% (31/39) of the Pre-ART samples along with two samples from the HIVNC and one from ART groups **(Figure 3b)**. Seven Pre-ART samples clustered with the HIVNC and ART groups. The HCA result was consistent with that of the MDS plot (**Supplementary Fig 1**) and principal component analysis.

**Figure 3.**
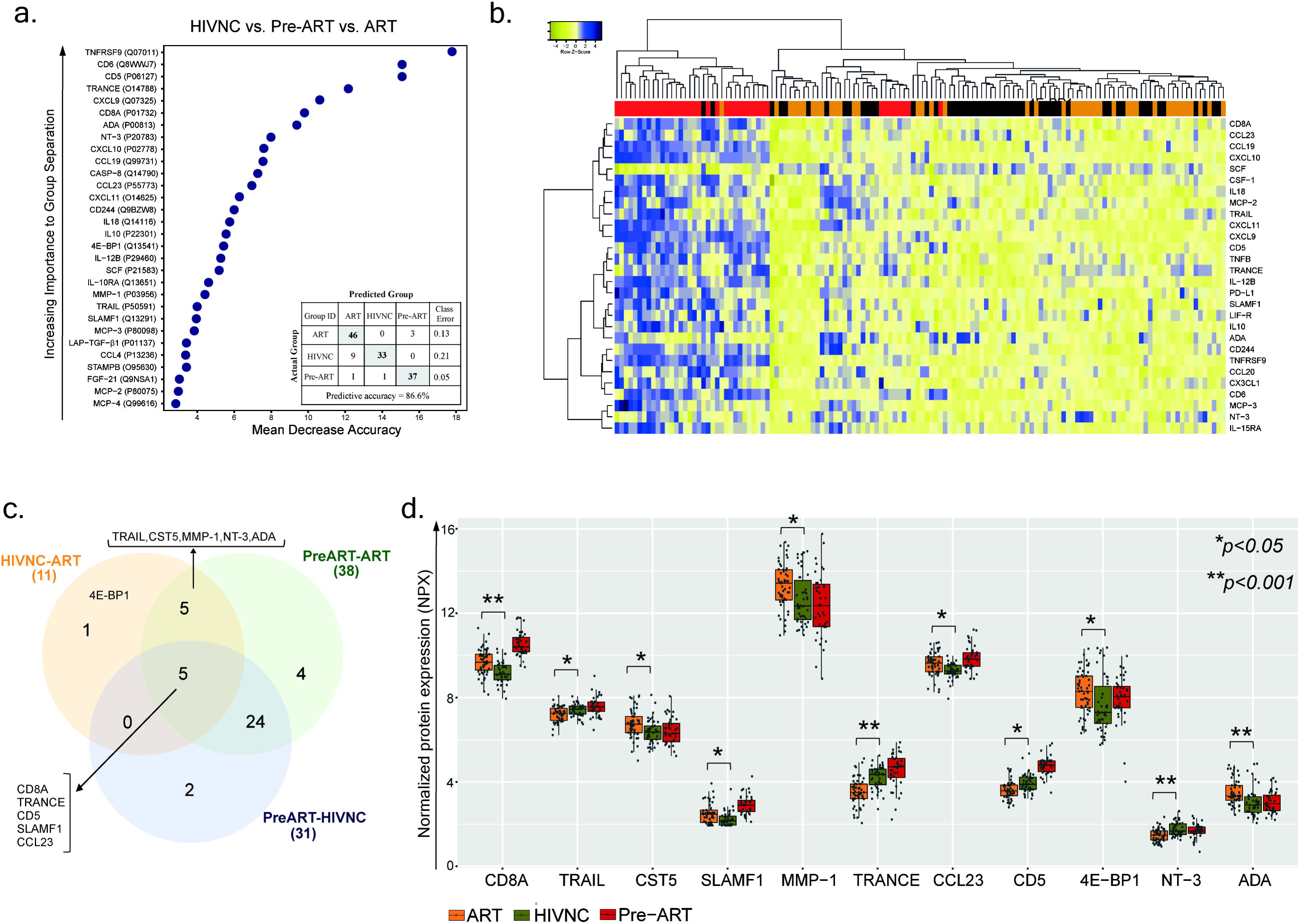
Plasma inflammation markers: (a) The random forest (RF) analysis of soluble factors resulted in predictive accuracies of 86.6% for HIVNC, Pre-ART and ART. The soluble factors importance plots displays top 30 metabolites which most strongly contribute to the groups’ separation for HIVNC, Pre-ART and ART. (b) Hierarchical clustering analysis of ANOVA of differentially expressed proteins with false discovery rate (FDR) <0.001, identified clustering of 79% (31/39) of Pre-ART samples along with two samples from the HIVNC and one from the ART group. The ART and HIVNC samples clustered separately from Pre-ART samples but intermingled with each other. (c) Venn diagram showing significantly different protein in the study group. The sum of the numbers in each large circle represents the total number of differentially expressed proteins in plasma in the different groups (HIVNC vs. ART, Pre-ART vs.ART and Pre-ART vs. HIVNC). The overlapping part of the circles represents significantly different proteins in the indicated groups. (d) Comparative analysis of 11 soluble markers that are significantly different between ART and HIVNC.

Among the 75 proteins, 41 showed a significant difference (*p*<0.05, Tukey HSD) between at least one of the groups compared (ART vs Pre-ART, Pre-ART vs HIVNC and ART vs HIVNC) (**Supplimentary table 1**). Levels of 31 inflammatory proteins were found to be significantly different between the Pre-ART and HIVNC groups, 38 proteins were significantly different between the Pre-ART and ART groups, and 11 proteins were significantly different (*p*<0.01, Tukey HSD) between the HIVNC and ART groups (**Figure 3c**). Out of 11 proteins, 4E-BP1 was found to differ significantly only between the ART and HIVNC groups, and not between any of the other groups. Five proteins were different between all the three groups (CD8A, TRANCE, sCD5, SLAMF1, and CCL23) (**Figure 3c**). Of the proteins that showed significant difference between the ART and HIVNC groups, levels of soluble NT3, CD5, TRAIL, and TRANCE were lower, but levels of ADA, MMP1, CST5 and 4E-BP1 were higher, when compared to both HIVNC and Pre-ART groups. On the other hand, CD8A, SLAMF1 and CCL23 were higher in the ART group as compared to the HIVNC group (**Figure 3d**).

### Telomere Length

Telomere length was analyzed only in two groups (HIVNC and ART) as PBMC were not available for the Pre-ART group. Linear regression analysis, after adjusting for chronological age and gender showed a significant negative association of HIV-1 positive status on telomere length (−2.84, 95%CI −4.012, −1.67, p<0.0001) with the ART group having significantly shorter telomeres. However, no significant association was observed between duration of treatment and telomere length after adjusting for age and markers of disease progression within the ART group. Several of the biomarkers considered were significantly associated with telomere length after adjustment for age, HIV status, and gender, these include CXCL1 and CD40 which are both weakly positively association with telomere length, MMP-10 and CX3CL1, which are positively associated with telomere length and OSM which is negatively associated; results are outlined in detail in supplementary Table 2. Within the ART group, after adjusting for age, gender, duration of treatment, HIV-1 reservoir, CD4 count at initiation, CD8/CD4 ratio and sCD14, we observed CXCL1 and TGF-α to have a significant association with increased telomere length (0.2905, 95%CI: 0.0029,0.5780 p=0.048) and (0.7865, 95%CI:0.1003,1.4727, p=0.026), respectively. We also found that IL-10RA was significantly associated with decreased telomere length (−1.79, 95%CI: −3.51, −0.07 p=0.042)

## Discussion

This study examined a cohort of PLHIV on long-term successful cART from India, and found that despite a median duration of eight years of suppressive cART, several soluble inflammatory markers that were found to be evaluated in HIV-1 infected untreated subjects, were also significantly elevated in the ART group as compared to the HIVNC group. This study found higher levels of the inflammatory protein CST5, a first in PLHIV, as compared to HIVNC, in addition to higher levels of 4E-BP1, SLAMF1, CCL23 and lower levels of NT3 proteins, a first in PLHIV on long term cART. This is the first study from India in standardized public health setting with standard first-line cART we observed that despite successful long-term cART, persistent immune activation and residual inflammation exist in PLHIV who are therefore at higher risk of inflamm-aging and age-related diseases.

It has been reported that certain age-related NICMs like diabetes mellitus, cardiovascular disease, cancer, bone fracture and renal failure, are more common among HIV-1 infected individuals as compared to the general population. Exposure to prolonged cART along with low grade inflammation and persistent immune activation seen in HIV-infection is thought to be the cause for the increased risk of NICMs (Guaraldi et al., 2011; Sokoya et al., 2017). Chronic inflammatory conditions and persistent immune activation are believed to be the major drivers of aging physiology. Inflammatory biomarkers of aging and their association with co-morbid diseases has been studied well in many elderly populations. Proteins like 4E-BP1 and association of mTOR with aging and age-related diseases (Johnson et al., 2013; Oka et al., 2017), has already been documented. However, there are no studies that have examined the response of plasma 4E-BP1 to ART. We found higher levels of 4E-BP1 in the ART group as compared to both HIVNC and the Pre-ART group, suggesting this marker may be associated with duration of HIV infection and/or exposure to anti-retroviral drugs.

In the case of well-studied immune activation markers, the ART group had significantly higher levels of sCD14 like that seen in the Pre-ART group. Contradictory results that have been reported from different studies on the effect of cART on sCD14 (Sandler et al., 2011; Hattab et al., 2015; van den Dries et al., 2017) due to a variety of contributors (Mudd and Brenchley, 2016; Dandona et al., 2017). On the other hand, sCD163, a marker of vascular inflammation (Subramanian et al., 2012) and neurocognitive impairment (Burdo et al., 2013), was higher in PLHIV than in healthy controls, which is in contrast to previous reports (Castley et al., 2014). The novelty of our study was to explore of a large panel of inflammatory markers and show significantly higher levels of MMP1 (Huang et al., 2012), ADA (Climent et al., 2009), CD8A (Nishanian et al., 1991), SLAMF1 (Farina et al., 2004) and CCL23 (Nardelli et al., 1999) in PLHIV on long term ART. Many of these markers have been reported to be linked with early stages of various age-related diseases.

sCD5 and TRAIL have been reported as biomarkers of inflammation and related diseases in both HIV-infected as well as uninfected individuals (Herbeuval et al., 2005; Balestrieri et al., 2007; Volpato et al., 2011). We found lower levels of sCD5 and TRAIL in the ART group as compared to the HIVNC group. Of particular mention is the association between lower plasma and cerebrospinal fluid levels of TRAIL and early Alzheimer Disease (AD) (Wu et al., 2015). CD5, a negative regulator of antigen receptor signal transduction in lymphocytes (Perez-Villar et al., 1999), is reported to stimulate the production of the anti-inflammatory cytokine IL-10 by B-cells (Gary-Gouy et al., 2002). Low levels of sCD5 in HIV-infected individuals, associated with the existing pro-inflammatory state could promote the development of age-related cancers and other diseases in these individuals. An earlier study conducted in a Swedish cohort reported normalization of sCD5 to a healthy state upon anti-HIV treatment (Sperk et al., 2018), which is in contrast to our findings. However, the Swedish cohort was on two decades long successful therapy and were initiated on ART in the early stages of HIV-infection, unlike our cohort which was started on ART only in stage 2 or 3 of the disease as per national guidelines and policies available then.

TRANCE/RANKL, an important regulator of bone metabolism, was found to be lower in the ART group than in HIVNC. There is existing literature to show that HIV-infected individuals treated with NRTIs and NNRTIs have lower than normal levels of circulating RANKL (Mora et al., 2007). Lower levels of TRANCE have been reported as an independent predictor of non-traumatic fracture indicating its effect on osteoclastogenesis (Schett et al., 2004). Though Cystatin D (CST5) has not been well studied in HIV, our study showed altered levels of this protein in HIV-1 infected individuals. To the best of our knowledge this is the first study reporting elevated levels of CST5 in HIV infection.

NT3, another inflammatory marker, is reported to be strongly associated with neurocognitive impairment in PLHIV (Bergman et al., 2000; Meeker et al., 2011) and MMP1 with senescence associated conditions (Benanti et al., 2002; Lei et al., 2016). Higher levels of these two proteins could be due to a favoring age-associated changes and premature aging in HIV-infected individuals with prolonged exposure to cART. The decreased expression of NT3 was also observed in colorectal cancer (Genevois et al., 2013), while increased levels of plasma MMP1 have been associated with coronary atherosclerosis (Castillo et al., 2010) and cancers (Warnecke-Eberz et al., 2016; Huang et al., 2018) further provide evidence to support the role of these proteins in disease development during aging.

Higher levels of CCL23 and SLAMF1, which are involved in monocyte activation, in the ART group could contribute to sustained release of sCD14 (Theil et al., 2005). Further, continued immune activation could be due to several other mechanisms including, sustained HIV-replication in reservoir sites like lymph nodes and gastrointestinal tract (Chun et al., 2005), low levels of HIV and its proteins (Meier et al., 2007; Anand et al., 2018), altered gut microbiota in PLHIV (Sessa et al., 2019) and different ART regimens (Hileman and Funderburg, 2017).

Telomere length (TL) is a marker of replicative senescence and is a well-known predictor of health outcome in aging populations. Chronic inflammation can potentially contribute to age-related diseases through increased production of reactive oxygen species that damage telomeres and lead to cellular and immunological senescence (Jurk et al., 2014). Several studies have noticed an association between TL and age-related diseases (Rizvi et al., 2014). In HIV infection, immune activation and expanded proliferation of leukocyte subsets immediately after infection, along with uncontrolled viremia have been associated with a rapid decrease in TL (Pathai et al., 2013), followed by increase in TL with ART initiation. However, telomere induced replicative senscence with long term ART has also been reported (Cobos Jimenez et al., 2016). Our study also found shorter telomeres in PLHIV on ART for >5 years and noticed a significant association between TL and inflammatory markers like CXCL1, TGF-α and IL10RA. Telomere length shortening has been well reported in several inflammatory conditions and related diseases like cancer. Zadka et al (2018) showed a positive correlation between IL10RA expression and disease pathogenesis in colorectal cancer (Zadka et al., 2018).

Our study has some limitations that merit comment. First, the HIV-infected population selected for this study were from the best pool of successfully treated individuals (free of co-infections and co-morbidities), through the Indian National ART program. The findings may not generalize to the general population of treated HIV-1 positive individuals. Second, due to the lack of earlier studies in this setting, the design of our study, and limited sample size, the conclusions drawn are limited to associations with modest significance. Also, many the statistical tests were run, but only the most significant results have been highlighted, for this reason, the results should be considered hypothesis generating. Third, the patients were not monitored virologically as a standard of care, and we only have virological data at the time of sampling. Any potential viral blips which are most unlikely reflected by the patients CD4 history could have introduced bias in the expression of inflammatory markers. However, among the few studies involving inflammatory markers in virally suppressed populations from LMICs (Margolick et al., 2017; Margolick et al., 2018), our study is the first comprehensive study on inflammation and aging, that compared a large panel of inflammatory markers in PLHIV on long term suppressive ART from India, which uses a standardized public health approach for treatment monitoring. Finally, the telomere length comparison between the ART group and the HIVNC could not be adjusted for health factors or disease progression markers, such as CD4 as these were not measured on HIVNC. As Telomere length was taken as the average for PBMCs, the different in CD4 and CD8 cells percentages may have contributed to the magnitude of the difference; further studies investigating difference based on cell type are needed.

In conclusion, we observed several soluble inflammatory biomarkers that differed significantly between PLHIV and healthy individuals, despite eight years of successful therapy-Interestingly, several of these biomarkers have been previously shown to be associated with inflammatory conditions like cancer, cardiovascular, neurological and skeletal diseases. Put together, these data suggest that HIV-1 infected individuals, even those on long term successful ART, may be at higher risk of developing inflammatory diseases leading to inflamm-aging, a low-grade chronic inflammation which is a major risk factor for the development of many age-associated diseases and to some extent morbidity and mortality in elderly people in the general population. In addition, we found a large difference in telomere length between subjects on long-term ART and HIVNC subjects, even after adjusting of age; this warrants further investigation.

## Acknowledgments

H.B. acknowledge support from the HIV Research Trust, UK supported in part by ViiV Health Care, Council of Scientific and Industrial Research (CSIR), India. Authors would like to thank technical support received from Dr. Ponnuraj CP, Mrs. Gomathi, Mr. Kannan Muthuramalingam, and Mr. Sathya Murthi.

## Contribution

H.B., S.S.A, N.R.M, M.S., N.C. performed the laboratory experiments; A.T.A. and E.E.G. performed bioinformatics and statistical analysis; U.N. and A.T.A. made the figures. N.P., R.S., V.J., and S.K.T. recruited study subjects and provided the clinical data. P.N. provided the clinical interpretation. U.N. and L.E.H conceived and designed the study; U.N. wrote the first draft of the paper reviewed by E.E.G, H.B., M.S., L.E.H. All the authors approved the final version of the manuscript.

