## Supplementary material for "Systemic inflammation and risk of age-associated diseases in people living with HIV on long term suppressive antiretroviral therapy": Supplimentary Table 1 and 2

**Supplementary Table 1.** Comparative analysis of the ANOVA significant proteins

|  |  | **Median** | | | **Level of Significance** | | |
| --- | --- | --- | --- | --- | --- | --- | --- |
| **Protein** | **ANOVA** | **ART** | **HIVNC** | **PreART** | **HIVNC vs ART** | **PreART vs. ART** | **PreART vs. HIVNC** |
| CD8A | <0.001 | 9.67 | 9.12 | 10.4 | <0.001 | <0.001 | <0.001 |
| MCP-3 | <0.001 | 1.42 | 1.36 | 1.97 | NS | <0.001 | <0.001 |
| CDCP1 | 0.047 | 3.16 | 2.97 | 3.29 | NS | NS | 0.043 |
| CD244 | <0.001 | 6.14 | 6.2 | 7.03 | NS | <0.001 | <0.001 |
| LAP-TGF-β1 | 0.031 | 6.25 | 6.19 | 6.56 | NS | 0.027 | NS |
| IL6 | 0.006 | 2.73 | 2.79 | 3.3 | NS | 0.009 | 0.021 |
| CXCL11 | <0.001 | 8.29 | 7.38 | 9.8 | NS | <0.001 | <0.001 |
| TRAIL | <0.001 | 7.25 | 7.45 | 7.56 | 0.038 | <0.001 | NS |
| CXCL9 | <0.001 | 7.08 | 6.84 | 9.33 | NS | <0.001 | <0.001 |
| CST5 | 0.006 | 6.74 | 6.37 | 6.31 | 0.013 | 0.024 | NS |
| CD6 | <0.001 | 4.84 | 5 | 6.75 | NS | <0.001 | 0 |
| SCF | <0.001 | 9.29 | 9.25 | 8.55 | NS | <0.001 | <0.001 |
| IL18 | <0.001 | 8.15 | 7.71 | 8.85 | NS | <0.001 | <0.001 |
| SLAMF1 | <0.001 | 2.47 | 2.16 | 2.9 | 0.018 | <0.001 | <0.001 |
| IL-10RA | 0.002 | -0.12 | -0.05 | 0.16 | NS | 0.002 | NS |
| MMP-1 | 0.002 | 13.42 | 12.33 | 12.36 | 0.033 | 0.002 | NS |
| LIF-R | <0.001 | 2.24 | 2.15 | 2.43 | NS | <0.001 | <0.001 |
| CCL19 | <0.001 | 8.51 | 8.69 | 10.16 | NS | <0.001 | <0.001 |
| IL-15RA | <0.001 | 0.28 | 0.26 | 0.46 | NS | <0.001 | <0.001 |
| PD-L1 | <0.001 | 4.35 | 4.28 | 4.77 | NS | <0.001 | <0.001 |
| TRANCE | <0.001 | 3.5 | 4.35 | 4.72 | <0.001 | <0.001 | 0.044 |
| IL-12B | <0.001 | 4.37 | 4.35 | 5.53 | NS | <0.001 | <0.001 |
| IL10 | <0.001 | 2.46 | 2.42 | 3.23 | NS | <0.001 | <0.001 |
| CCL23 | <0.001 | 9.63 | 9.25 | 9.83 | 0.004 | 0.021 | <0.001 |
| CD5 | <0.001 | 3.57 | 3.88 | 4.79 | 0.001 | <0.001 | <0.001 |
| CCL3 | 0.003 | 4.63 | 4.5 | 4.99 | NS | 0.037 | 0.003 |
| CXCL10 | <0.001 | 8.97 | 9.01 | 11.26 | NS | <0.001 | <0.001 |
| 4E-BP1 | 0.038 | 8.29 | 7.29 | 8.04 | 0.03 | NS | NS |
| DNER | 0.009 | 8.47 | 8.6 | 8.35 | NS | 0.026 | 0.013 |
| CD40 | 0.035 | 10.36 | 10.29 | 10.76 | NS | 0.026 | NS |
| FGF-19 | 0.017 | 7.08 | 7.62 | 7.93 | NS | 0.024 | NS |
| MCP-2 | <0.001 | 7.63 | 7.41 | 8.15 | NS | 0.021 | <0.001 |
| CASP-8 | 0.007 | 2.79 | 2.38 | 3.24 | NS | NS | 0.006 |
| CX3CL1 | <0.001 | 5.34 | 5.38 | 5.61 | NS | <0.001 | <0.001 |
| TNFRSF9 | <0.001 | 5.76 | 5.74 | 7.27 | NS | <0.001 | <0.001 |
| NT-3 | <0.001 | 1.48 | 1.67 | 1.67 | <0.001 | 0.012 | NS |
| TWEAK | 0.018 | 9.17 | 9.11 | 9.1 | NS | 0.048 | 0.023 |
| CCL20 | <0.001 | 5.69 | 5.59 | 6.26 | NS | <0.001 | 0.005 |
| ADA | <0.001 | 3.33 | 2.91 | 2.96 | <0.001 | <0.001 | NS |
| TNFB | <0.001 | 3.65 | 3.77 | 4.28 | NS | <0.001 | <0.001 |
| CSF-1 | <0.001 | 8.77 | 8.77 | 8.99 | NS | <0.001 | <0.001 |

NS- Non-significant

**Supplementary Table 2.** Proteins with significant association with Telomere length after adjustment for HIV status, age and gender in ART and HIVNC

| Protein | Estimated difference | 95% CI | P-value |
| --- | --- | --- | --- |
| CXCL1 | 0.3921 | (0.002, 0.782) | 0.0486 |
| MMP-10 | 1.7882 | (0.936, 0.640) | 0.0001 |
| CD40 | 0.7412 | (0.029, 1.454) | 0.0417 |
| CX3CL1 | 1.4148 | (0.200, 2.630) | 0.0232 |
| OSM | -0.7886 | (-1.403, -0.174) | 0.0127 |
