## Supplementary material for "Systemic inflammation and risk of age-associated diseases in people living with HIV on long term suppressive antiretroviral therapy": Supplimentary Fig 1

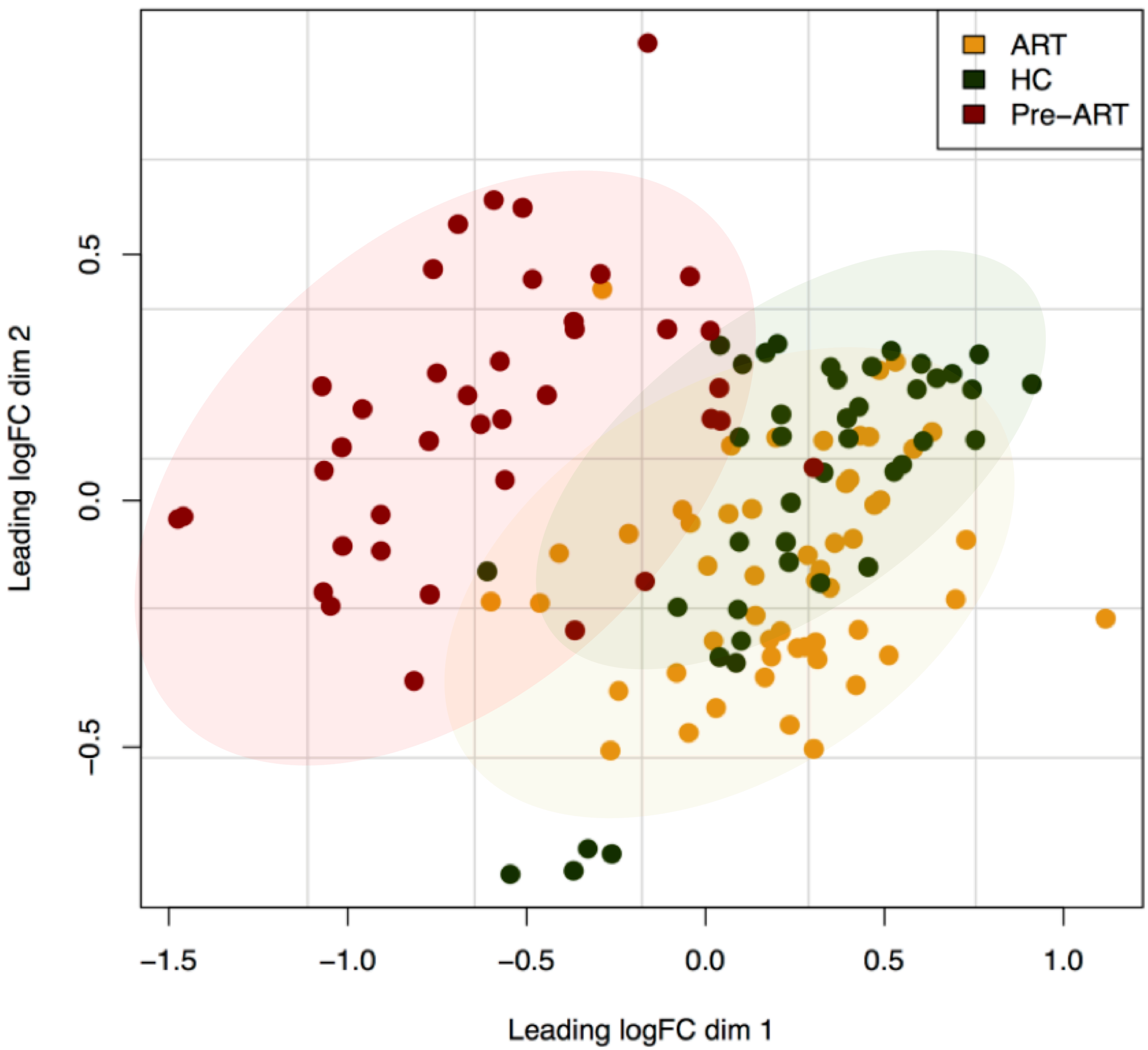


**Supplementary figure 1**. Multi-dimensional scale (MDS) plot for clustering of patient samples based on inflammatory protein level.
